# GWAS meta-analysis highlights the hypothalamic-pituitary-gonadal axis (HPG axis) in the genetic regulation of menstrual cycle length

**DOI:** 10.1101/333708

**Authors:** Triin Laisk, Viktorija Kukuškina, Duncan Palmer, Samantha Laber, Chia-Yen Chen, Teresa Ferreira, Nilufer Rahmioglu, Krina Zondervan, Christian Becker, Jordan W. Smoller, Margaret Lippincott, Andres Salumets, Ingrid Granne, Stephanie Seminara, Benjamin Neale, Reedik Mägi, Cecilia M. Lindgren

**Affiliations:** Department of Obstetrics and Gynecology, Institute of Clinical Medicine, University of Tartu, Tartu, Estonia; Estonian Genome Center, Institute of Genomics, University of Tartu, Tartu, Estonia; Competence Centre on Health Technologies, Tartu, Estonia; Analytic and Translational Genetics Unit, Department of Medicine, Massachusetts General Hospital and Harvard Medical School, Boston, Massachusetts, USA; Stanley Center for Psychiatric Research, Broad Institute of Harvard and MIT, Cambridge, Massachusetts, USA; Big Data Institute, Li Ka Shing Center for Health for Health Information and Discovery, Oxford University, Oxford, UK; Wellcome Centre for Human Genetics, University of Oxford, Oxford, UK; Psychiatric and Neurodevelopmental Genetics Unit, Massachusetts General Hospital, Boston, Massachusetts, USA; Broad Institute of MIT and Harvard, Boston, Massachusetts, USA; Oxford Endometriosis CaRe Centre, Nuffield Department of Women’s and Reproductive Health, University of Oxford, UK; Harvard Reproductive Sciences Center and Reproductive Endocrine Unit, Massachusetts General Hospital, Boston, Massachusetts, USA; Department of Obstetrics and Gynecology, University of Helsinki and Helsinki University Hospital, Helsinki, Finland; Department of Biomedicine, Institute of Biomedicine and Translational Medicine, University of Tartu, Tartu, Estonia; Nuffield Department of Women’s and Reproductive Health, University of Oxford, UK; Program in Medical and Population Genetics, Broad Institute, Boston, MA, USA

## Abstract

The normal menstrual cycle requires a delicate interplay between the hypothalamus, pituitary, and ovary. Therefore, its length is an important indicator of female reproductive health. Menstrual cycle length has been shown to be partially controlled by genetic factors, especially in the follicle stimulating hormone beta-subunit (*FSHB*) locus. GWAS meta-analysis of menstrual cycle length in 44,871 women of European ancestry confirmed the previously observed association with the *FSHB* locus and identified four additional novel signals in, or near, the *GNRH1, PGR, NR5A2* and *INS-IGF2* genes. These findings confirm the role of the hypothalamic-pituitary-gonadal axis in the genetic regulation of menstrual cycle length, but also highlight potential novel local regulatory mechanisms, such as those mediated by *IGF2*.

## Introduction

A menstrual cycle is crucial for human reproduction as it is required for oocyte selection, maturation, and ovulation in preparation for its fertilization and subsequent pregnancy [1]. The median menstrual cycle length is 27-30 days, depending on age [2] and can be divided into two distinct ovarian phases - the follicular and luteal phases separated by ovulation. During the follicular phase the emerging follicle secretes estrogen that causes proliferation of the endometrium, the uterine lining, and in the subsequent luteal phase progesterone secretion from the corpus luteum of the ruptured follicle causes endometrium to cease proliferating and change both phenotypically and functionally in preparation for implantation of the embryo [3]. The menstrual cycle and its length are under the control of reproductive hormones secreted via the integration of the hypothalamic–pituitary–gonadal axis (HPG axis), where the gonadotropin releasing hormone (GnRH) secreted from the hypothalamus stimulates the release of the gonadotropins, follicle stimulating hormone (FSH) and luteinizing hormone (LH), from the anterior pituitary [3,4]. FSH and LH in turn stimulate follicular growth and secretion of estrogens to prepare for ovulation, and progesterone from ovarian follicular cells [3,4]. The length of menstrual cycle reflects fertility status, and has been associated with a range of reproductive traits, such as time to pregnancy, risk of spontaneous abortion, and success rates in assisted reproduction [5–7]. Moreover, shorter cycles have been associated with an increased risk of a gynecological condition known as endometriosis [8]. Although a small twin study suggested no significant heritability for menstrual cycle length [9], it was recently demonstrated that a genetic variant in the promoter of follicle stimulating hormone beta subunit gene (*FSHB*) is associated with longer menstrual cycles, nulliparity and lower endometriosis risk [PMID: 26732621]. However, only variants in or near the *FSHB* gene reached genome-wide significance among 9,534 women [10], leaving the possibility that additional loci regulating menstrual cycle length could be revealed in larger studies.

Here, we present the results of a genome-wide association study (GWAS) meta-analysis of 44,871 women of European ancestry. We confirm the previous association with the *FSHB* locus [10] and also identify a further four novel association signals, contributing to an increase in our knowledge on the underlying genetics of menstrual cycle length control along the hypothalamus-pituitary-ovarian axis, and also providing a genetic basis for the observed epidemiological correlations with gynecological pathologies.

## Results

### Genome-wide association signals for menstrual cycle length

A total of five loci reached genome-wide significance (linear regression P < 5×10^−8^) for association with menstrual cycle length in the meta-analysis including data from two cohorts and a total of 44,871 women (**Table 1, Figure 1, Supplementary Figure 1**). The strongest signal (rs11031006, P_meta_ = 3.6 ×; 10^−36^,*β* _UKBB_ = −0.16 (s.e. = 0.01)) is in strong LD (r^2^ = 0.80) with the previously reported variant in *FSHB* promoter (rs10835638), while the remaining four loci are signals previously not reported. The strongest novel association (rs6670899, P_meta_ = 6.6×10^−13^, _β UKBB_ = −0.06 (s.e. = 0.01)) is 105 kb upstream of the *NR5A2* gene, which encodes a DNA-binding zinc finger transcription factor that is implicated in regulation of steroidogenesis during granulosa cell differentiation [11]. This same region has previously been associated with age at menarche [12] (lead signal rs6427782 A-allele (r^2^ = 0.45 with rs6670899) was shown to increase age at menarche [12], and increases menstrual cycle length in our analysis, P_meta_ = 4.7 ×; 10^−6^). second intron of the *DOCK5* gene, but in strong LD (r^2^ = 0.90) with rs6185 (P_meta_ = 2.0 ×; 10^−10^), a The second novel signal (rs13261573, P_meta_ = 1.2 ×; 10^−10^, *β*_UKBB_ = −0.07 (s.e. = 0.01)) is in the missense variant in the gonadotropin releasing hormone 1 gene (*GNRH1*). *GNRH1* encodes the precursor for a peptide in the gonadotropin releasing hormone family that regulate the release of FSH and LH from the anterior pituitary [3,4]. We also observed two additional signals on chromosome 11; the first (lead signal rs471811, P_meta_ = 3.0 ×;10^−8^, *β* _UKBB_ = −0.03 (s.e. = 0.01)) lies 42kb upstream of progesterone receptor gene (*PGR*) and 14 kb downstream of a PGR antisense RNA (*PGR-AS1*). The second novel signal on chromosome 11 (rs11042596, P_meta_ = 4.5 ×; 10^−8^,*β* _UKBB_ = 0.04 (s.e. = 0.01)), is located 31 kb downstream the *INS-IGF2* and *IGF2* genes.

**Table 1.** Genetic variants associated with menstrual cycle length

| Region | Nearest gene(s) | SNP | Alleles<br>Other allele/<br>Effect allele<br>(EAF) | UKBB |  | EGCUT |  | Meta-analysis |
| --- | --- | --- | --- | --- | --- | --- | --- | --- |
|  |  |  |  | Effect<br>(SD of the<br>binned<br>menstrual<br>cycle<br>length) | P-value | Effect<br>(SD of the<br>binned<br>menstrual<br>cycle<br>length) | P-value | P-value |
| 11:30226528 | *FSHB* | rs11031006 | A/G (0.86) | -0.16 (0.01) | $1.1 \times 10^{-38}$ | -0.06 (0.02) | $6.6 \times 10^{-4}$ | $3.6 \times 10^{-36}$ |
| 1:199891438 | *NR5A2* | rs6670899 | A/C (0.57) | -0.05 (0.01) | $1.1 \times 10^{-10}$ | -0.04 (0.01) | $4.7 \times 10^{-4}$ | $6.6 \times 10^{-13}$ |
| 8:25248615 | *DOCK5/*<br>*GNRH1* | rs13261573 | A/G (0.75) | -0.07 (0.01) | $1.7 \times 10^{-11}$ | -0.02 (0.01) | $7.0 \times 10^{-2}$ | $1.2 \times 10^{-10}$ |
| 11:10104420<br>3 | *PGR/*<br>*PGR-AS1* | rs471811 | C/T (0.31) | -0.03 (0.01) | $4.8 \times 10^{-5}$ | -0.06 (0.01) | $6.3 \times 10^{-5}$ | $3.0 \times 10^{-8}$ |
| 11:2118860 | *IGF2/ INS-*<br>*IGF2* | rs11042596 | G/T (0.34) | 0.04 (0.01) | $1.1 \times 10^{-7}$ | 0.02 (0.01) | $3.5 \times 10^{-2}$ | $4.5 \times 10^{-8}$ |

**Figure 1.**
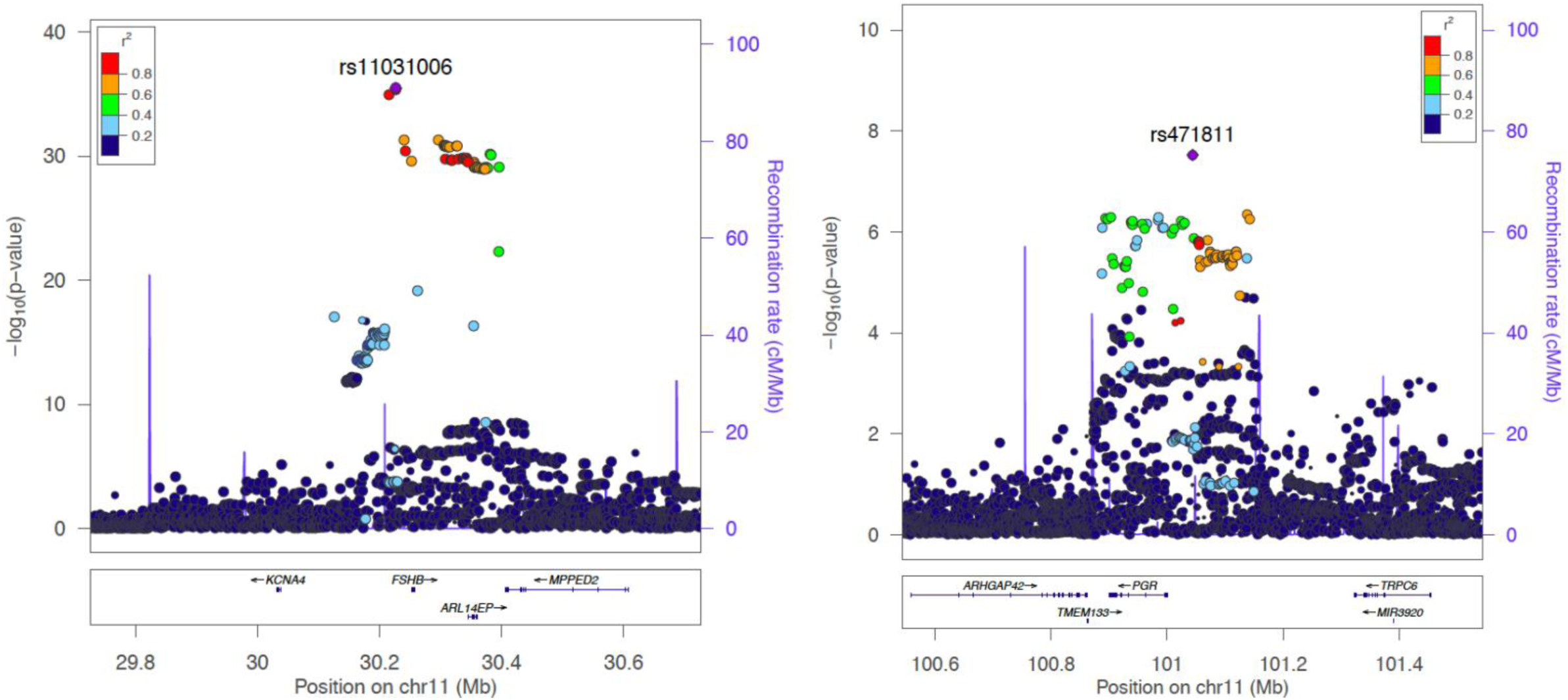
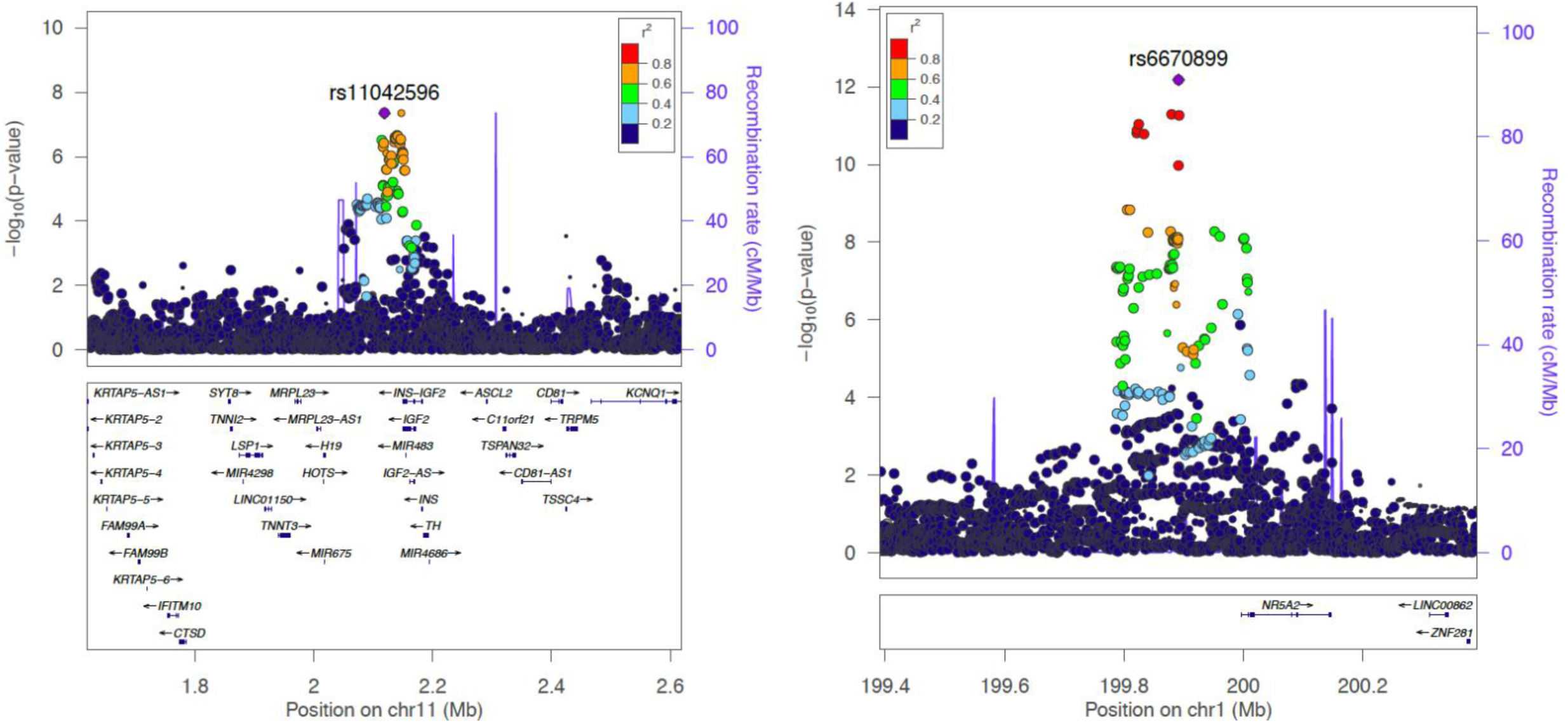
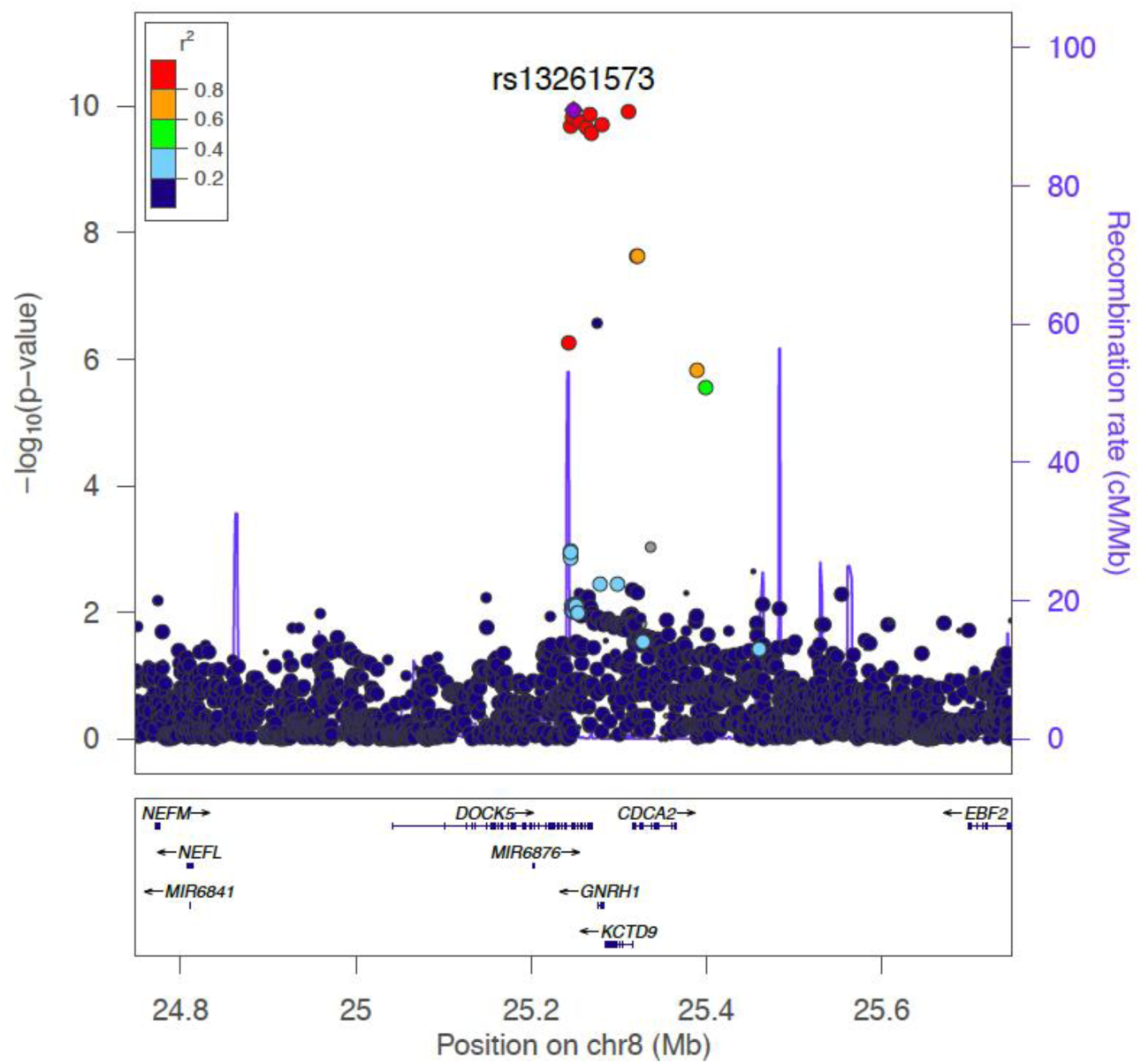
Regional plots for five genome-wide significant loci. Regional plot depicts SNPs plotted by their position and GWAS meta-analysis – log10(P-value) for association with menstrual cycle length. Nearby genes are shown on the lower panel.

### SNP-based heritability of menstrual cycle length and menstrual cycle irregularities

We evaluated SNP-based heritability (phenotypic variance explained by SNPs in the GWAS meta-analysis) using LDSC (LD-score regression) [13]. The overall SNP-based heritability of menstrual cycle length was estimated at 6.1% (s.e. = 1.2). After filtering out all variants within± 500kb of the lead SNPs, the heritability estimate for menstrual cycle length decreased to 5.4% (s.e. = 1.1) indicating that common SNPs explain a small but significant part of menstrual cycle length variability, and moreover, the majority of the SNP-heritability still remains to be discovered.

### Gene based associations of menstrual cycle length and irregular menstruation

A MAGMA (Multi-marker Analysis of GenoMic Annotation; [14]) genome-wide gene association analysis of our GWAS meta-analysis summary statistics highlighted ten genes that passed the suggested threshold for significance (P = 2.7 ×; 10^−6^, Bonferroni correction for association testing of 18,297 protein coding genes); *ARL14EP, SMAD3, MPPED2, RHBDD1, IGF2, COL4A4, PGR, INS-IGF2, FSHB*, and *ARHGEF3* (**Supplementary Table 1**). Six of these genes (*ARL14EP/FSHB/MPPED2, IGF2, INS-IGF2, PGR*) overlap with three loci identified in the single marker analysis, while the remaining four novel gene signals did not harbor genome-wide significant SNPs (lowest P-values for SNPs in *SMAD3, RHBDD1, COL4A4,* and *ARHGEF3* were rs11856909 P = 6.2 ×;10^−8^; rs4673173 P = 1.0 ×; 10^−7^, rs12467261 P = 1.3 ×;10^−7^, and rs73086331 P = 1.9 ×;-610 respectively).

### Genetic associations between menstrual cycle length and other traits

To evaluate the potential shared genetic architecture between menstrual cycle length and other traits, we performed a look-up in the GWAS catalogue (https://www.ebi.ac.uk/gwas/; **Supplementary Table 2**) for menstrual cycle length associated variants and candidate SNPs identified by FUMA. Several significant associations were found for the *FSHB* locus, including gonadotropin (FSH and LH) levels, age at menarche and menopause, spontaneous dizygotic twinning, endometriosis and PCOS (P <= 3 ×; 10^−8^). Additionally, the *NR5A2* locus was associated with menarche timing (P = 5 ×; 10^−8^), and showed some evidence for association with age at voice drop (P = 6 ×; 10^−7^) and pancreatic cancer (P = 1 ×; 10^−11^).

Next, to determine whether other phenotypes were associated with loci regulating menstrual cycle length, we conducted a PheWAS using the sentinel markers for each locus (rs11031006, rs6670899, rs13261573, rs471811, rs11042596) and the UKBB phenotypes present in the Oxford BIG browser [http://big.stats.ox.ac.uk/]. Associations with a P < 2.1 × 10^−5^ (corresponding to a Bonferroni-corrected threshold of 0.05/2,419) are shown in **Supplementary Table 3.** Again, the *FSHB* locus (rs11031006) showed the largest number of associations, including three genome-wide significant associations (P < 5 × 10^−8^) with “Years since last cervical smear”, “Bilateral oophorectomy (both ovaries removed)”, “Diagnoses - main ICD10: N92 Excessive, frequent and irregular menstruation” in UKBB. Nominally significant associations were also observed for “Age when periods started (menarche)” (P = 1.7 ×; 10^−7^), “Non-cancer illness code, self-reported: endometriosis” (P = 3.8 ×; 10^−7^), “Part of a multiple birth” (P = 4.6×10^−7^), supporting the findings from the GWAS catalogue look-up. In this comparison, the allele associated with longer cycles decreased the risk of oophorectomy, menstrual cycle disturbances and endometriosis, and was associated with later menarche. Similarly, rs6670899 (*NR5A2*) menstrual cycle lengthening allele was associated with later menarche timing (P = 3.2 ×; 10^−8^).

Finally, we carried out a genetic correlation analysis with the LDSC method implemented in LD-Hub [15]. Comparison with cardiometabolic, anthropometric, autoimmune, hormone, cancer and reproductive traits (for example lowest P-values were observed for age of first birth (r_g_ = 0.12, s.e. = 0.07, P = 0.055) and age at menopause (r_g_ = 0.15, s.e. = 0.08, P = 0.058)) revealed no significant correlations (**Supplementary Table 4**).

### Functional annotation of associated variants and candidate gene mapping

Functional mapping and annotation of genetic associations for menstrual cycle length was carried out using FUMA [16], and a total of 600 candidate SNPs (defined as being in LD with the lead SNPs with a r^2^ >= 0.6) were identified. The majority of these (∼90%; **Supplementary Figure 2A** and **Supplementary Table 5**) were located in intergenic or intronic regions, and >75% of the variants overlapped chromatin state annotations (**Supplementary Figure 2C**; **Supplementary Table 5)**, suggesting that they affect gene regulation.

To identify the potential effector transcripts for the five significant loci for menstrual cycle length, genes within the loci were prioritized if there was evidence for both eQTL and chromatin interaction [16].

In the *FSHB* locus, a total of two lead SNPs (rs11031006 and rs11032051), eight independent (r^2^< 0.6) significant SNPs and 359 candidate SNPs were identified (**Supplementary Table 5**). Numerous significant eQTL associations (FDR <0.05) were identified in different datasets (**Supplementary Table 6**), but genes that were highlighted by both eQTL and chromatin interaction mapping included *FSHB, ARL14EP* and *MPPED2* (**Supplementary Figure 3**).

The *INS-IGF2* locus (lead signal rs11042596) included a total of 34 candidate SNPs, with the lowest RDB score (1d - likely to affect binding and linked to expression of a gene target) for rs6578986. eQTL mapping and chromatin interactions highlighted *IGF2* and *INS-IGF2* as likely effector transcripts at this locus (**Supplementary Table 6, Supplementary Figure 3**).

The *PGR* locus (lead signal rs471811) included a total of 61 candidate SNPs, and *ANGPTL5* was prioritized by both eQTL (thyroid in GTEx_v7, [17]; FDR < 0.05) and chromatin interaction analysis.

In the *NR5A2* locus on chromosome 1 (lead variant rs6670899), four independent significant SNPs and 133 candidate SNPs were identified (**Supplementary Table 5**), six of which have evidence for likely affecting regulatory element binding (RDB score 2; **Supplementary Table 5)**. Two genes were prioritized based on eQTL data (*ZNF281* in dorsolateral prefrontal cortex [18]; and C1orf106 in testis (GTEx_v7: [17]; FDR < 0.05), while *ZNF281* was also additionally mapped using chromatin interaction data.

Finally, in the *DOCK5-GnRH* locus on chromosome 8 (lead variant rs132661573), 13 potential candidate SNPs were identified **Supplementary Table 5)**, including rs6185, a missense variant in the *GnRH* gene. Seven of the 13 candidates are also eQTLs for *GnRH* in whole blood (GTEx_v7: [17], FDR < 0.05).

### Tissue specificity and gene set enrichment analysis

Using the list of genes that were highlighted either in gene-based analysis and/or had both eQTL and chromatin interaction data supporting their candidacy, we performed a tissue specificity and pathway enrichment analysis with the GENE2FUNC option implemented in FUMA [16]. Enrichment test of differentially expressed genes across GTEx_v7 30 tissue types (see **Methods**) showed significantly higher expression of prioritized genes in female reproductive tissues: uterus (Bonferroni corrected P-value; P_Bon_ = 0.047), cervix uteri (P_Bon_ = ^0.048), and ovary (PBon = 0.050) (Supplementary Figure 4; Supplementary Table 7). Prioritized^ genes were also overrepresented in hormone activity related pathways (for example, GO hormone activity FDR = 7.6 ×;^−7^-^3^10, KEGG GnRH signaling pathway FDR = 1.5 ×; 10, WikiPathways [19] ovarian infertility genes FDR = 7.5 ×; 10^−5^ (**Supplementary Table 8**). Tissue and cell-type enrichment analysis with DEPICT [20] revealed no significant enrichments.

Using GREAT [21] we found that genes within the five significant menstrual cycle length GWAS loci are enriched for uterus and circulating hormone level-related mouse phenotypes (**Supplementary Table 9**) and further highlighted an enrichment at these loci for “genes involved in hormone ligand-binding receptors” (P_FDR_ = 1.3 ×; 10^−2^; **Supplementary Table 10**). Reviewing the MGI mouse phenotype database [22] showed that mouse knockouts of *Fshb, Nr5a2, Gnrh1,* and *Pgr* all present with female reproductive phenotypes (**Supplementary Table 11**), including altered estrous cycle length or abnormal ovulation for *Fshb, Gnrh1,* and *Pgr* (progesterone receptor) (**Supplementary Table 12**). *Nr5a2* (nuclear receptor subfamily 5, group A, member 2) is linked to reduced fertility, primarily by reduced circulating progesterone levels in *Nr5a*2+/−female mice [23].

The presence of female reproductive phenotypes in mice with altered expression of *Fshb, Nr5a2, Gnrh1,* and *Pgr* provides evidence that these genes may be causal and could explain, at least in part, the mediating mechanisms underlying four out of the five significant loci associated with menstrual cycle length. Further experimental validation will be necessary to fully unravel the mechanism of these non-coding associations.

## Discussion

This large-scale GWAS meta-analysis reveals several novel insights into the genetic control of menstrual cycle length and provides evidence of the genetic underpinnings of the epidemiological associations between menstrual cycle length and other traits. Understanding the genetics regulating normal menstrual cycle variation is vital for figuring out the mechanisms leading to different menstrual cycle-related pathologies. Moreover, genetic control of menstrual cycle and folliculogenesis is of importance for *in vitro* fertilization treatment, where markers allowing for individualization of treatment protocols are still extensively sought [24].

While some of the results confirm what is already known about the biology of the menstrual cycle (such as the regulatory role of GnRH and FSH in the HPG axis), others point to potentially novel modulators and the role of local control of folliculogenesis. For example, IGF2 has been proposed to be an important local regulator of folliculogenesis [25], as it stimulates estrogen production [26] and modulates the action of FSH and LH, whereas IGF2 expression in turn is regulated by FSH [27]. However, to our knowledge no direct link between genetic variation in the *INS-IGF2* region and menstrual cycle length had been previously demonstrated. Similarly, while it is known that progesterone is the dominant hormone in the second half of the menstrual cycle, the evidence linking genetic variation in the progesterone signaling pathway with menstrual cycle length was scarce [28,29]. SMAD3, highlighted in gene-based analysis, is shown to modulate the proliferation of follicular granulosa cells and also ovarian steroidogenesis [30], and is an essential regulator of FSH signaling in the mouse [31]. Recently, genetic variation in *SMAD3* was associated with dizygotic twinning [32]. However, the obvious candidacy and support for one gene in most of these loci does not exclude the possibility that there might be additional genes and/or functional sequence in these loci that contribute to menstrual cycle length.

Analysis of pleiotropy between menstrual cycle length associated variants and GWAS signals of other traits confirmed the central role of the *FSHB* locus, which is involved in regulating the reproductive lifespan from menarche to menopause, and is also associated with gynecological diseases such as polycystic ovary syndrome (PCOS) and endometriosis, and with menstrual cycle disturbances. While there is epidemiological evidence that shorter menstrual cycles are associated with earlier age at menopause [33], we did not observe a significant overlap on a genetic level, as these traits did not show a significant genetic correlation. At the same time, the *FSHB* locus is significantly associated with both menstrual cycle length and age at menopause [10,34], indicating that this locus is probably largely driving the observed phenotypic correlation between menstrual cycle length and age at menopause in the literature. Our study has a number of limitations. The UKBB includes women over 40 years of age, some of whom are approaching menopause and might therefore have more irregular and shorter cycles [35]. Also, participants in the UKBB were asked about their *current* cycle length, whereas EGCUT participants were asked to report their cycle length at the age of 25-35, where it is believed to be most regular [35]. Although it is possible that the effect estimates from these two cohorts may not be directly comparable, we observe consistency in effect direction and magnitude. Finally, while our sample size is the largest to date, it may still be underpowered to detect further associations.

In conclusion, the largest menstrual cycle length GWAS meta-analysis to date confirms the role of key players in the HPG axis in the genetic regulation of menstrual cycle length (*GNRH1, FSHB, PGR*) but also pinpoints novel genes with a potential local regulatory role (such as *IGF2*/*INS-IGF2* and *NR5A2*). Our analysis also highlights the central role of the *FSHB* locus in female reproductive health and provides evidence that the systemic determinants of normal menstrual cycle length (*FSHB*) are also associated with menstrual cycle-related pathologies, such as excessive, frequent and irregular menstruation. However, the loci identified as significant in our analysis represent a small fraction of the SNP-heritability for menstrual cycle length, warranting additional larger meta-analysis efforts to further uncover the remaining genetic underpinnings of menstrual cycle length.

## Methods

### Study cohorts

The current meta-analysis included a total of 44,871 women of European ancestry from two cohorts. We used the data of the UK Biobank (UKBB), a population-based biobank comprising 502,637 people (aged 37-73) recruited from across the UK during 2006-2010, who have filled out detailed medical history questionnaires [36]. Menstrual cycle length information was derived from data field 3710 “Length of menstrual cycle”. Participants were asked “How many days is your usual menstrual cycle? (The number of days between each menstrual period)”. This question was asked of women who had indicated they were not menopausal and still had menstrual periods in their answer to data field 2724 (“Have you had your menopause (periods stopped)?”). The phenotype was transformed according to the default PHESANT pipeline [37], whereby the integer phenotype is split into three ordered bins if a single value represents >20% of all respondents answers. As a result, length of menstrual cycle was split into <26, 26-28, and >= 28 days. All answers corresponding to “Irregular cycle”, “Do not know”, and “Prefer not to answer” were coded as NA. As a result, each bin included 14,211 (mean age 45.7 years, range 39-69), 4,949 (45.7 years (40-70)), and 29,227 (45.9 years (40-70)) individuals, respectively. Additionally, individuals were filteredas described in https://github.com/Nealelab/UK_Biobank_GWAS, leaving 30,245 individuals for final analysis.

We also included data from the Estonian Biobank (EGCUT), a population-based biobank with 51,515 participants of European ancestry [38]. In EGCUT, women older than 35 were asked about their menstrual cycle length using the question: “Approximately how long was your menstrual cycle when you were between 25 and 35 years old?”, with the following choices: I don’t know; I have not had any menstrual cycles; Irregular; 20 days or less; 21-24 days; 25-29 days; 30-35 days; more than 35 days. To follow a similar structure as with the UKBB data, the answers were regrouped into three bins: <25; 25-29; >=30 days, resulting in 2,877 (56.3 years (33-95)), 10,354 (54.3 years (33-101)), and 1,395 (50.9 years (34-96)) individuals in each bin, respectively.

### GWAS and meta-analysis

In the UKBB dataset, quality control and association testing were carried out as described in https://github.com/Nealelab/UK_Biobank_GWAS. In brief, samples were filtered for white British genetic ancestry, related individuals, individuals with sex chromosome aneuploidies, and individuals who had withdrawn their participation in the UKBB. The analysis included SNPs imputed to the Haplotype Reference Consortium (HRC) reference panel, additional filters included minor allele frequency (MAF) > 0.1%, HWE P > 1×10^−10^, and imputation INFO score > 0.8. Association testing was carried out using linear regression implemented in HAIL [https://github.com/hail-is/hail], adjusting for the first 10 principal components (PCs).

In EGCUT, Illumina Human CoreExome, OmniExpress, 370CNV BeadChip and GSA arrays were used for genotyping. Quality control included filtering on the basis of sample call rate (< 98%), heterozygosity (> mean ± 3SD), genotype and phenotype sex discordance, cryptic relatedness (IBD > 20%) and outliers from the European descent based on the MDS plot in comparison with HapMap reference samples. SNP quality filtering included call rate (<99%), MAF (<1%) and extreme deviation from Hardy–Weinberg equilibrium (P < 1 ×;10^−4^). Imputation was performed using SHAPEIT2 for prephasing, the Estonian-specific reference panel [PMID: 28401899] and IMPUTE2 [PMID: 19543373] with default parameters. Association testing was carried out with EPACTS [https://github.com/statgen/EPACTS], adjusting for 10 PCs and age at recruitment.

Before meta-analysis, results from individual cohorts underwent central quality control with EasyQC [39], checking for allele frequency against the HRC reference, and filtering out variants with a MAF < 1% and INFO score < 0.4. The results from individual cohorts were meta-analysed with METAL [40] using sample-size weighted P-value-based meta-analysis with genomic control correction. The meta-analysis included 9,344,826 markers, and those with a P < 5 ×; 10^−8^ were considered genome-wide significant.

To convert the effects obtained from the linear regression of binned trait to a standardized scale, we calculated the mean and variance of the 0, 1, 2 binned menstrual cycle length phenotype, and divided the effect estimates from linear regression with calculated standard deviation of the binned phenotype.

### Gene-based testing

Gene-based genome-wide association analysis was carried out with MAGMA 1.6 [14] with default settings implemented in FUMA [16]. Briefly, variants were assigned to protein-coding genes (n = 18,297; Ensembl build 85) if they are located in the gene body, and the resulting SNP P-values are combined into a gene test-statistic using the SNP-wise mean model [14]. According to the number of tested genes, the level of genome wide significance was set at 0.05/18,297 = 2.7 ×; 10^−6^.

### Heritability estimate

The menstrual cycle length GWAS meta-analysis summary statistics and LD Score Regression (LDSC) method [13] were used for heritability estimation. The linkage disequilibrium (LD) estimates from European ancestry samples in the 1000 Genomes Project were used as a reference.

### Functional mapping

Functional annotation was performed using the FUMA platform designed for prioritisation, annotation and interpretation of GWAS results [16]. As the first step, independent significant SNPs in the GWAS meta-analysis summary statistics were identified based on their P-value (P < 5 ×; 10^−8^) and independence from each other (r^2^ < 0.6 in the 1000G phase 3 reference) within a 1Mb window. Thereafter, lead SNPs were identified from independent significant SNPs, which are independent of each other (r^2^ < 0.1). SNPs that were in LD with the identified independent SNPs (r^2^ ≥ 0.6) within a 1Mb window, have a MAF of ≥ 1% and GWAS meta-analysis P-value > 0.05, were selected as candidate SNPs and taken forward for further annotation.

FUMA annotates candidate SNPs in genomic risk loci based on functional consequences on genes (ANNOVAR; [41]), CADD (a continuous score showing how deleterious the SNP is to protein structure/function; scores greater than 12.37 indicate potential pathogenicity [42]) and RegulomeDB scores (ranging from 1-7, where lower score indicates greater evidence for having regulatory function) [43], 15 chromatin states from the Roadmap Epigenomics Project [44,45], eQTL data (GTEx v6 and v7 [17]), blood eQTL browser [46], BIOS QTL browser [47], BRAINEAC [48], MuTHER [49], xQTLServer [50], and the CommonMind Consortium [18]), and 3D chromatin interactions from HI-C experiments of 21 tissues/cell types [51], also embedded in the FUMA platform. Next, genes were mapped using positional mapping, which is based on ANNOVAR annotations and maximum distance between SNPs (default 10kb) and genes, eQTL mapping, and chromatin interaction mapping. Chromatin interaction mapping was performed with significant chromatin interactions (defined as FDR < 1 ×; 10^−6^). The two ends of significant chromatin interactions were defined as follows: region 1 – a region overlapping with one of the candidate SNPs, and region 2 – another end of the significant interaction, used to map to genes based on overlap with a promoter region (250bp upstream and 50bp downstream of the transcription start site).

### Genetic associations between menstrual cycle length and other traits

The Oxford Brain Imaging Genetics (BIG) Server (v2.0) [http://big.stats.ox.ac.uk/] was used to query the sentinel variants in each locus against an array of UKBB phenotypes (**Supplementary Table 3**). Additionally, during the FUMA functional mapping, sentinel SNPs and proximal SNPs in tight LD(r^2^ = 0.6) were linked with the GWAS catalog [https://www.ebi.ac.uk/gwas/]. Full results of the GWAS catalog query are shown in **Supplementary Table 2**.

We analysed genome-wide genetic correlation analyses applying the LDSC method [13] using the LD-Hub resource and 50 selected traits (cardiometabolic, anthropometric, autoimmune, hormone, reproductive, cancer, and aging categories). Full results of the LDSC genetic correlation analysis are reported in **Supplementary Table 4.**

### Tissue specificity and gene set enrichment analyses

Tissue and gene set enrichment analyses were carried out with GENE2FUNC implemented in FUMA [16]. Genes that were highlighted in MAGMA gene-based analysis or which had functional annotation support from eQTL and chromatin interaction data were used as an input (a total of 14 genes). 2 ×; 2 enrichment tests were carried out, using all genes as a background gene set. The GTEx v7 30 general tissue types dataset was used for tissue specificity analyses. Differentially expressed gene (DEG) sets are pre-calculated in the GENE2FUNC by performing two-sided t-test for any one of tissues against all others. For this, expression values were normalized (zero-mean) following a log_2_ transformation of expression values (transcripts per million - TPM). Genes with P ≤ 0.05 after Bonferroni correction and absolute log fold change ≥ 0.58 were defined as differentially expressed genes in a given tissue compared to others. In addition to general DEG, upregulated and downregulated DEG sets were also pre-calculated by taking sign of t-statistics into account. Our set of prioritized input genes was tested against each of the DEG sets using a hypergeometric test, where background genes are genes that have average expression value > 1 in at least one of the tissues. Significant enrichment at Bonferroni corrected P ≤ 0.05 are coloured in red in **Supplementary Figure 4.**

### Tools used in this paper

HAIL: https://github.com/hail-is/hail

Oxford BIG browser: http://big.stats.ox.ac.uk/ FUMA: http://fuma.ctglab.nl/

GWAS catalog: https://www.ebi.ac.uk/gwas/

GREAT: http://great.stanford.edu/public/html/

HaploReg: http://archive.broadinstitute.org/mammals/haploreg/haploreg.php

MGI: http://www.informatics.jax.org/phenotypes.shtml

## Acknowledgements

The research in this paper has been carried out using the UK Biobank resource (application 17085). The work has received funding from the European Commission Horizon 2020 research and innovation programme under grant agreement 692065 (project WIDENLIFE), grants IUT34-16 and PUTJD726 from the Estonian Ministry of Education and Research, grant EU49695 from Enterprise Estonia and grant 5P50HD028138-27 (Common Complex Trait Genetics of Reproductive Phenotypes) from NIH/NICHD. S.L. has a Postdoctoral Research Fellowship funded by Novo Nordisk. T.F. is supported by the NIHR Biomedical Research Centre, Oxford. C.M.L is supported by the Li Ka Shing Foundation, WT-SSI/John Fell funds and by the NIHR Biomedical Research Centre, Oxford, by Widenlife and NIH (5P50HD028138-27).

## Supporting information

**S1 Fig. Manhattan and QQ plot for menstrual cycle length GWAS meta-analysis.**

**S2 Fig. Functional annotation for all SNPs with r^2^**≥**0.6 with the sentinel SNPs**.

a) Percentage of SNPs according to their functional category; b) Percentage of SNPs according to their RegulomeDB score. Lower score indicates a more likely regulatory role; c) Percentage of SNPs according to their minimum chromatin state across 127 tissues. Lower score indicates a more likely regulatory role.

**S3. Fig. Circos plots demonstrating the results of eQTL and chromatin interaction mapping for loci on chromosome 11, 1, and 8, respectively.**

The outermost layer depicts SNPs with a P < 0.05 in a Manhattan-style plot. SNPs in detected genomic risk locus are coloured according to their r^2^ value to the most significant variant at the locus. Y-axis values range from 0 to the maximum –log10(P-value) of the SNPs. The second and third layer represent the chromosome ring, where the risk locus is coloured in blue, and genes mapped by chromatin interactions with this region, depicted by orange ribbons, are displayed. eQTL associations are shown by green ribbons, and genes that were mapped by both eQTL associations and chromatin interaction data are highlighted in red.

**S4 Fig. Expression of prioritized genes and results of differential expression analysis.**

a) Heatmap displaying average expression of prioritized genes and b) differential expression analysis of prioritized genes across GTEx v7 30 general tissue types.

**S1 Table. Results of the MAGMA gene-based analysis.**

**S2 Table. Results of the GWAS catalog lookup S3 Table. Associated phenotypes in UKBB**

**S4 Table. Results of the genetic correlation analysis performed using the LD-Score method S5 Table. Candidate SNPs outlined by FUMA analysis and functional annotation**

**S6 Table. eQTL look-up results from FUMA analysis**

**S7 Table. Tissue specificity analysis from GENE2FUNC implemented in FUMA**

**S8 Table. Geneset analysis results from GENE2FUNC implemented in FUMA**

**S9 Table. GREAT analysis for enriched mouse phenotypes**

**S10 Table. GREAT MSigDB pathway enrichment analysis**

**S11 Table. Mouse phenotypes of genes prioritized by FUMA and GREAT for the 5 menstrual cycle length GWAS meta-analysis signals**

**S12 Table. Mouse female reproductive phenotypes of genes prioritized by FUMA and GREAT for the 5 menstrual cycle length GWAS meta-analysis signals**

## References

1. Jabbour HN, Kelly RW, Fraser HM, Critchley HOD. Endocrine regulation of menstruation. Endocr Rev. 2006;27: 17–46. doi:10.1210/er.2004-0021

2. Guo Y, Manatunga AK, Chen S, Marcus M. Modeling menstrual cycle length using a mixture distribution. Biostatistics. 2006;7: 100–14. doi:10.1093/biostatistics/kxi043

3. Reed BG, Carr BR. The Normal Menstrual Cycle and the Control of Ovulation. Endotext. 2000.

4. Barbieri RL. The endocrinology of the menstrual cycle. Methods Mol Biol. 2014;1154: 145–69. doi:10.1007/978-1-4939-0659-8_7

5. Small CM, Manatunga AK, Klein M, Feigelson HS, Dominguez CE, McChesney R, et al. Menstrual cycle characteristics: associations with fertility and spontaneous abortion. Epidemiology. 2006;17: 52–60.

6. Wise LA, Mikkelsen EM, Rothman KJ, Riis AH, Sørensen HT, Huybrechts KF, et al. A prospective cohort study of menstrual characteristics and time to pregnancy. Am J Epidemiol. 2011;174: 701–9. doi:10.1093/aje/kwr130

7. Brodin T, Bergh T, Berglund L, Hadziosmanovic N, Holte J. Menstrual cycle length is an age-independent marker of female fertility: results from 6271 treatment cycles of in vitro fertilization. Fertil Steril. 2008;90: 1656–61. doi:10.1016/j.fertnstert.2007.09.036

8. Matalliotakis IM, Cakmak H, Fragouli YG, Goumenou AG, Mahutte NG, Arici A. Epidemiological characteristics in women with and without endometriosis in the Yale series. Arch Gynecol Obstet. 2008;277: 389–93. doi:10.1007/s00404-007-0479-1

9. van den Akker OB, Stein GS, Neale MC, Murray RM. Genetic and environmental variation in menstrual cycle: histories of two British twin samples. Acta Genet Med Gemellol (Roma). 1987;36: 541–8.

10. Ruth KS, Beaumont RN, Tyrrell J, Jones SE, Tuke MA, Yaghootkar H, et al. Genetic evidence that lower circulating FSH levels lengthen menstrual cycle, increase age at menopause and impact female reproductive health. Hum Reprod. 2016; doi:10.1093/humrep/dev318

11. Saxena D, Escamilla-Hernandez R, Little-Ihrig L, Zeleznik AJ. Liver receptor homolog-1 and steroidogenic factor-1 have similar actions on rat granulosa cell steroidogenesis. Endocrinology. 2007;148: 726–34. doi:10.1210/en.2006-0108

12. Perry JRB, Day F, Elks CE, Sulem P, Thompson DJ, Ferreira T, et al. Parent-of-origin-specific allelic associations among 106 genomic loci for age at menarche. Nature. 2014;514: 92–7. doi:10.1038/nature13545

13. Bulik-Sullivan B, Finucane HK, Anttila V, Gusev A, Day FR, Loh P-R, et al. An atlas of genetic correlations across human diseases and traits. Nat Genet. 2015;47: 1236–41. doi:10.1038/ng.3406

14. de Leeuw CA, Mooij JM, Heskes T, Posthuma D. MAGMA: generalized gene-set analysis of GWAS data. Tang H, editor. PLoS Comput Biol. 2015;11: e1004219. doi:10.1371/journal.pcbi.1004219

15. Zheng J, Erzurumluoglu AM, Elsworth BL, Kemp JP, Howe L, Haycock PC, et al. LD Hub: a centralized database and web interface to perform LD score regression that maximizes the potential of summary level GWAS data for SNP heritability and genetic correlation analysis. Bioinformatics. 2016; btw613. doi:10.1093/bioinformatics/btw613

16. Watanabe K, Taskesen E, van Bochoven A, Posthuma D. Functional mapping and annotation of genetic associations with FUMA. Nat Commun. 2017;8: 1826. doi:10.1038/s41467-017-01261-5

17. Lonsdale J, Thomas J, Salvatore M, Phillips R, Lo E, Shad S, et al. The Genotype-Tissue Expression (GTEx) project. Nat Genet. Nature Research; 2013;45: 580–585. doi:10.1038/ng.2653

18. Fromer M, Roussos P, Sieberts SK, Johnson JS, Kavanagh DH, Perumal TM, et al. Gene expression elucidates functional impact of polygenic risk for schizophrenia. Nat Neurosci. 2016;19: 1442–1453. doi:10.1038/nn.4399

19. Kelder T, van Iersel MP, Hanspers K, Kutmon M, Conklin BR, Evelo CT, et al. WikiPathways: building research communities on biological pathways. Nucleic Acids Res. 2012;40: D1301–7. doi:10.1093/nar/gkr1074

20. Pers TH, Karjalainen JM, Chan Y, Westra H-J, Wood AR, Yang J, et al. Biological interpretation of genome-wide association studies using predicted gene functions. Nat Commun. 2015;6: 5890. doi:10.1038/ncomms6890

21. McLean CY, Bristor D, Hiller M, Clarke SL, Schaar BT, Lowe CB, et al. GREAT improves functional interpretation of cis-regulatory regions. Nat Biotechnol. 2010;28: 495–501. doi:10.1038/nbt.1630

22. Blake JA, Eppig JT, Kadin JA, Richardson JE, Smith CL, Bult CJ, et al. Mouse Genome Database (MGD)-2017: community knowledge resource for the laboratory mouse. Nucleic Acids Res. 2017;45: D723–D729. doi:10.1093/nar/gkw1040

23. Labelle-Dumais C, Paré J-F, Bélanger L, Farookhi R, Dufort D. Impaired progesterone production in Nr5a2+/− mice leads to a reduction in female reproductive function. Biol Reprod. 2007;77: 217–25. doi:10.1095/biolreprod.106.059121

24. Altmäe S, Hovatta O, Stavreus-Evers A, Salumets A. Genetic predictors of controlled ovarian hyperstimulation: where do we stand today? Hum Reprod Update. 2011;17: 813–28. doi:10.1093/humupd/dmr034

25. el-Roeiy A, Chen X, Roberts VJ, LeRoith D, Roberts CT, Yen SS. Expression of insulin-like growth factor-I (IGF-I) and IGF-II and the IGF-I, IGF-II, and insulin receptor genes and localization of the gene products in the human ovary. J Clin Endocrinol Metab. 1993;77: 1411–8. doi:10.1210/jcem.77.5.8077342

26. Spicer LJ, Aad PY. Insulin-like growth factor (IGF) 2 stimulates steroidogenesis and mitosis of bovine granulosa cells through the IGF1 receptor: role of follicle-stimulating hormone and IGF2 receptor. Biol Reprod. 2007;77: 18–27. doi:10.1095/biolreprod.106.058230

27. Baumgarten SC, Convissar SM, Zamah AM, Fierro MA, Winston NJ, Scoccia B, et al. FSH Regulates IGF-2 Expression in Human Granulosa Cells in an AKT-Dependent Manner. J Clin Endocrinol Metab. 2015;100: E1046–55. doi:10.1210/jc.2015-1504

28. Taylor KC, Small CM, Epstein MP, Sherman SL, Tang W, Wilson MM, et al. Associations of progesterone receptor polymorphisms with age at menarche and menstrual cycle length. Horm Res Paediatr. 2010;74: 421–7. doi:10.1159/000316961

29. Rowe EJ, Eisenstein TK, Meissler J, Rockwell LC. Gene x environment interactions impact endometrial function and the menstrual cycle: PROGINS, life history, anthropometry, and physical activity. Am J Hum Biol. 2013;25: 681–94. doi:10.1002/ajhb.22430

30. Liu Y, Chen X, Xue X, Shen C, Shi C, Dong J, et al. Effects of Smad3 on the proliferation and steroidogenesis in human ovarian luteinized granulosa cells. IUBMB Life. 2014;66: 424–37. doi:10.1002/iub.1280

31. Gong X, McGee EA. Smad3 is required for normal follicular follicle-stimulating hormone responsiveness in the mouse. Biol Reprod. 2009;81: 730–8. doi:10.1095/biolreprod.108.070086

32. Mbarek H, Steinberg S, Nyholt DR, Gordon SD, Miller MB, McRae AF, et al. Identification of Common Genetic Variants Influencing Spontaneous Dizygotic Twinning and Female Fertility. Am J Hum Genet. 2016;98: 898–908. doi:10.1016/j.ajhg.2016.03.008

33. Whelan EA, Sandler DP, McConnaughey DR, Weinberg CR. Menstrual and reproductive characteristics and age at natural menopause. Am J Epidemiol. 1990;131: 625–32.

34. Stolk L, Perry JRB, Chasman DI, He C, Mangino M, Sulem P, et al. Meta-analyses identify 13 loci associated with age at menopause and highlight DNA repair and immune pathways. Nat Genet. 2012;44: 260–8. doi:10.1038/ng.1051

35. Mihm M, Gangooly S, Muttukrishna S. The normal menstrual cycle in women. Anim Reprod Sci. 2011;124: 229–236. doi:10.1016/j.anireprosci.2010.08.030

36. Sudlow C, Gallacher J, Allen N, Beral V, Burton P, Danesh J, et al. UK Biobank: An Open Access Resource for Identifying the Causes of a Wide Range of Complex Diseases of Middle and Old Age. PLoS Med. 2015;12: e1001779. doi:10.1371/journal.pmed.1001779

37. Millard LAC, Davies NM, Gaunt TR, Davey Smith G, Tilling K. Software Application Profile: PHESANT: a tool for performing automated phenome scans in UK Biobank. Int J Epidemiol. 2017;47: 29–35. doi:10.1093/ije/dyx204

38. Leitsalu L, Haller T, Esko T, Tammesoo M-L, Alavere H, Snieder H, et al. Cohort Profile: Estonian Biobank of the Estonian Genome Center, University of Tartu. Int J Epidemiol. 2015;44: 1137–1147. doi:10.1093/ije/dyt268

39. Winkler TW, Day FR, Croteau-Chonka DC, Wood AR, Locke AE, Mägi R, et al. Quality control and conduct of genome-wide association meta-analyses. Nat Protoc. 2014;9: 1192–1212. doi:10.1038/nprot.2014.071

40. Willer CJ, Li Y, Abecasis GR. METAL: fast and efficient meta-analysis of genomewide association scans. Bioinformatics. 2010;26: 2190–2191. doi:10.1093/bioinformatics/btq340

41. Wang K, Li M, Hakonarson H. ANNOVAR: functional annotation of genetic variants from high-throughput sequencing data. Nucleic Acids Res. 2010;38: e164. doi:10.1093/nar/gkq603

42. Kircher M, Witten DM, Jain P, O’Roak BJ, Cooper GM, Shendure J. A general framework for estimating the relative pathogenicity of human genetic variants. Nat Genet. 2014;46: 310–5. doi:10.1038/ng.2892

43. Boyle AP, Hong EL, Hariharan M, Cheng Y, Schaub MA, Kasowski M, et al. Annotation of functional variation in personal genomes using RegulomeDB. Genome Res. 2012;22: 1790–1797. doi:10.1101/gr.137323.112

44. Dunham I, Kundaje A, Aldred SF, Collins PJ, Davis CA, Doyle F, et al. An integrated encyclopedia of DNA elements in the human genome. Nature. 2012;489: 57–74. doi:10.1038/nature11247

45. Roadmap Epigenomics Consortium A, Kundaje A, Meuleman W, Ernst J, Bilenky M, Yen A, et al. Integrative analysis of 111 reference human epigenomes. Nature. 2015;518: 317–30. doi:10.1038/nature14248

46. Westra H-J, Peters MJ, Esko T, Yaghootkar H, Schurmann C, Kettunen J, et al. Systematic identification of trans eQTLs as putative drivers of known disease associations. Nat Genet. 2013;45: 1238–1243. doi:10.1038/ng.2756

47. Zhernakova D V, Deelen P, Vermaat M, van Iterson M, van Galen M, Arindrarto W, et al. Identification of context-dependent expression quantitative trait loci in whole blood. Nat Genet. 2016;49: 139–145. doi:10.1038/ng.3737

48. Ramasamy A, Trabzuni D, Guelfi S, Varghese V, Smith C, Walker R, et al. Genetic variability in the regulation of gene expression in ten regions of the human brain. Nat Neurosci. 2014;17: 1418–1428. doi:10.1038/nn.3801

49. Grundberg E, Small KS, Hedman ÅK, Nica AC, Buil A, Keildson S, et al. Mapping cis- and trans-regulatory effects across multiple tissues in twins. Nat Genet. 2012;44: 1084–9. doi:10.1038/ng.2394

50. Ng B, White CC, Klein H-U, Sieberts SK, McCabe C, Patrick E, et al. An xQTL map integrates the genetic architecture of the human brain’s transcriptome and epigenome. Nat Neurosci. 2017;20: 1418–1426. doi:10.1038/nn.4632

51. Schmitt A, Hu M, Jung I, Xu Z, Qiu Y, Tan C, et al. A Compendium of Chromatin Contact Maps Reveals Spatially Active Regions in the Human Genome. Cell Rep. 2016;17: 2042–2059. doi:10.1016/j.celrep.2016.10.061

